# Locus-specific ChIP combined with NGS analysis reveals genomic regulatory regions that physically interact with the *Pax5* promoter in a chicken B cell line

**DOI:** 10.1101/089821

**Authors:** Toshitsugu Fujita, Fusako Kitaura, Miyuki Yuno, Yutaka Suzuki, Sumio Sugano, Hodaka Fujii

**Affiliations:** Chromatin Biochemistry Research Group, Combined Program on Microbiology and Immunology, Research Institute for Microbial Diseases, Osaka University, Suita, Osaka, Japan; Department of Medical Genome Sciences, Graduate School of Frontier Sciences, The University of Tokyo, Kashiwa, Chiba, Japan; Department of Medical Genome Sciences, Graduate School of Frontier Sciences, The University of Tokyo, Minato-ku, Tokyo, Japan

## Abstract

Chromosomal interactions regulate genome functions, such as transcription, via dynamic chromosomal organization in the nucleus. In this study, we identified genomic regions that physically bind to the promoter region of the *Pax5* gene in the chicken B-cell line DT40, with the goal of obtaining mechanistic insight into transcriptional regulation through chromosomal interaction. Using insertional chromatin immunoprecipitation (iChIP) in combination with next-generation sequencing (NGS) (iChIP-Seq), we found that the *Pax5* promoter bound to multiple genomic regions. The identified chromosomal interactions were independently confirmed by *in vitro* engineered DNA-binding molecule-mediated ChIP (*in vitro* enChIP) in combination with NGS (*in vitro* enChIP-Seq). Comparing chromosomal interactions in wild-type DT40 with those in a macrophage-like counterpart, we found that some of the identified chromosomal interactions were organized in a B cell–specific manner. In addition, deletion of a B cell–specific interacting genomic region in chromosome 11, which was marked by active enhancer histone modifications, resulted in moderate but significant down-regulation of *Pax5* transcription. Together, these results suggested that *Pax5* transcription in DT40 cells is regulated by inter-chromosomal interactions. Moreover, these analyses showed that iChIP-Seq and *in vitro* enChIP-Seq are useful for non-biased identification of functional genomic regions that physically interact with a locus of interest.

## Introduction

Elucidation of the molecular mechanisms underlying genome functions, such as transcription, requires identification of molecules that interact with the genomic regions of interest. To this end, we developed locus-specific chromatin immunoprecipitation (locus-specific ChIP) technology [see review (Fujita and Fujii 2013b; Fujii and Fujita 2015; Fujita and Fujii 2016)]. Locus-specific ChIP is a biochemical tool for specifically isolating genomic regions of interest from cells. In combination with downstream biochemical analyses, one can identify molecules that physically interact with target genomic regions in cells.

In principle, locus-specific ChIP consists of locus-tagging and affinity purification. On the basis of various strategies for locus-tagging, we developed two locus-specific ChIP technologies, insertional ChIP (iChIP) (Hoshino and Fujii 2009; Fujita and Fujii 2012a) and engineered DNA-binding molecule-mediated ChIP (enChIP) (Fujita and Fujii 2013a; Fujita et al. 2013). iChIP utilizes an exogenous DNA-binding protein, such as a bacterial protein LexA, and its binding element for locus-tagging, whereas enChIP employs engineered DNA-binding molecules, such as transcription activator–like (TAL) proteins (Moscou and Bogdanove 2009; Boch et al. 2009) and the clustered regularly interspaced short palindromic repeats (CRISPR) system (Jinek et al. 2012; Cong et al. 2013), for the same purpose. After isolation of tagged loci by affinity purification, their interacting molecules can be comprehensively identified by downstream analyses including mass spectrometry (MS), next-generation sequencing (NGS), and microarrays. In fact, we have successfully identified proteins that interact with target loci by iChIP or enChIP in combination with MS, including a quantitative form of MS, stable isotope labeling with amino acids in cell culture (SILAC) (iChIP/enChIP-MS or -SILAC) (Fujita and Fujii 2011a, 2013a; Fujita et al. 2013; Fujita and Fujii 2014b; Fujita et al. 2015a). In addition, identification of chromatin-binding RNAs is also feasible using enChIP in combination with RT-PCR (enChIP-RT-PCR) or RNA sequencing (enChIP-RNA-Seq) (Fujita et al. 2013, 2015b).

Genome functions are mediated by chromosomal interactions (e.g., interactions between enhancers and promoters). To detect physical chromosomal interactions, several techniques have been utilized to date, including fluorescence *in situ* hybridization (FISH) (Trask 1991) and chromosome conformation capture (3C) plus 3C-derived methods (Dekker 2002; Simonis et al. 2006; Dostie et al. 2006; Lieberman-aiden et al. 2009; Dekker et al. 2013). In this regard, locus-specific ChIP can also be applied to detection of physical chromosomal interactions (one-to-many interactions). In fact, using iChIP in combination with microarrays (iChIP-microarray), McCullagh et al. succeeded in non-biased identification of genomic regions that interact with a target locus in yeast (McCullagh et al. 2010). More recently, we showed that it is also feasible to analyze physical chromosomal interactions using enChIP combined with NGS analysis (enChIP-Seq) (Fujita et al. 2016b).

The *Pax5* gene encodes a transcription factor essential for B-cell lineage commitment (Cobaleda et al. 2007). Disruption of the *Pax5* gene inhibits B-cell differentiation (Urbánek et al. 1994; Nutt et al. 1997), and *Pax5*-deficient B cells can be trans-differentiated into other lymphoid cell types in mice (Nutt et al. 1999; Mikkola et al. 2002; Schaniel et al. 2002). To obtain mechanistic insight into transcriptional regulation of the *Pax5* gene, we previously used iChIP-SILAC to identify proteins that interact with the *Pax5* promoter region in the chicken B-cell line DT40 (Fujita et al. 2015a). However, the mechanisms underlying regulation of *Pax5* transcription by chromosomal interactions remain incompletely understood. Although intron 5 of the mouse *Pax5* gene contains enhancers essential for transcription of the gene (Decker et al. 2009), it remains unclear whether similar regulatory mechanisms exist across species. In this regard, because the DNA sequences of *Pax5* intron 5 are scarcely conserved between mouse and chicken, it is possible that transcription of *Pax5* is controlled in a species-specific manner.

In this study, we applied iChIP in combination with NGS analysis (iChIP-Seq) to direct identification of genomic regions that interact with the *Pax5* promoter region in DT40 cells. Some of the detected chromosomal interactions were independently confirmed by an updated form of enChIP-Seq. In addition, deletion of a B cell–specific interacting genomic region significantly decreased *Pax5* transcription in DT40 cells, suggesting that the deleted region is an enhancer and that *Pax5* transcription is regulated through chromosomal interactions between this enhancer and the promoter. Thus, locus-specific ChIP in combination with NGS analysis revealed a mechanism of transcriptional regulation of the chicken *Pax5* gene.

## Results and Discussion

### Scheme of iChIP-Seq for analysis of chromosomal interactions around the *Pax5* promoter region

The scheme of iChIP-Seq used in this study is as follows (Figure 1A): (I) Using homologous recombination, binding elements of the bacterial DNA-binding protein LexA (LexA BE) were inserted ~0.3 kb upstream from the transcription start site (TSS) of *Pax5* exon 1A in chromosome Z of DT40 cells. (II) 3xFNLDD, which consists of 3xFLAG-tag, a nuclear localization signal (NLS), and LexA DNA-binding and dimerization domains, was expressed in the cells established in Step (I). (III) The resultant cells were crosslinked with formaldehyde and lysed, and chromatin DNA was fragmented by sonication. (IV) The tagged locus (*Pax5* promoter region) was affinity-purified using an anti-FLAG antibody. (V) After reverse crosslinking and DNA purification, genomic regions interacting with the *Pax5* promoter region were identified by NGS analysis.

In this study, we utilized DT40-derived cell lines (Figure 1B), which were previously established for iChIP-SILAC analysis of the *Pax5* promoter region (Fujita et al. s2015a); Non-KI(B) is DT40 expressing 3xFNLDD, and KI(B) is a DT40-derived cell line harboring an insertion of LexA BE in the *Pax5* promoter region and expressing 3xFNLDD. In our previous study, insertion of LexA BE and expression of 3xFNLDD did not disturb transcription of the endogenous *Pax5* gene (Fujita et al. 2015a), suggesting that the regulatory machinery involved for *Pax5* transcription is retained in both Non-KI(B) and KI(B). In addition, we previously showed that the *Pax5* promoter region can be efficiently isolated from KI(B) by iChIP (~10% of input as DNA yields) (Fujita et al. 2015a). Following the experimental scheme (Figure 1A), we isolated the *Pax5* promoter region by iChIP and subjected the purified DNA samples to NGS analysis using a HiSeq. NGS reads corresponding to the *Pax5* promoter region were clearly enriched when iChIP was performed with KI(B) but not Non-KI(B) [iChIP(#1) in Figure 1C]. A biological replicate of the iChIP-Seq analysis showed similar results [iChIP(#2) in Figure 1C]. These results demonstrated efficient isolation of the *Pax5* promoter region by iChIP.

**Figure 1.**
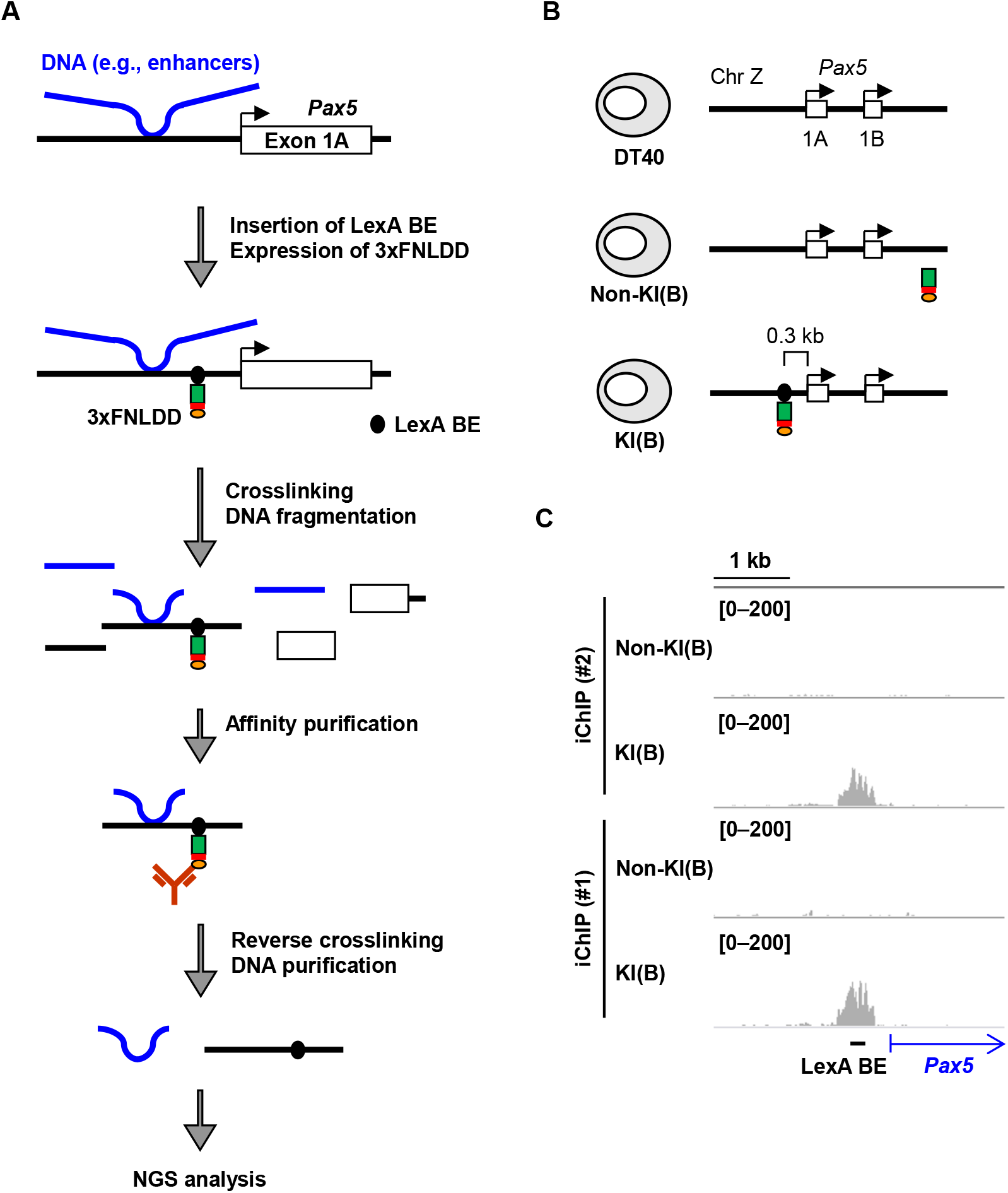
iChIP-Seq for identification of genomic regions that physically interact with the *Pax5* promoter in chicken B cells. (**A**) A schematic diagram of iChIP-Seq in this study. First, LexA-binding elements (LexA BE) were inserted into the *Pax5* promoter region, and 3xFNLDD [a fusion protein of the 3xFLAG-tag, a nuclear localization signal (NLS), and LexA DNA-binding domain plus dimerization domain] was expressed in DT40. After crosslinking with formaldehyde, chromatin DNA was fragmented by sonication, and the target locus was affinity-purified with anti-FLAG antibody. After reversal of crosslinking, DNA was purified and subjected to NGS analysis. (**B**) The chicken B-cell line DT40 and its derivatives used for iChIP-Seq. Non-KI(B): DT40 expressing 3xFNLDD, KI(B): DT40 containing LexA BE in the *Pax5* promoter region and expressing 3xFNLDD. The LexA BE was inserted 0.3 kb upstream from the transcription start site of *Pax5* exon 1A. KI: Knock-In. (**C**) Images of NGS peaks around the *Pax5* promoter region. NGS data from iChIP-Seq were visualized in IGV. The vertical viewing range (*y*-axis shown as Scale) was set to 0–200 based on the magnitude of the noise peaks.

### Detection of genomic regions that physically interact with the *Pax5* promoter region in DT40

Next, we proceeded to identify the genomic regions that interact with the *Pax5* promoter region in DT40 (Figure 2). Because 3xFNLDD might interact with endogenous DNA sequences, similar to the recognition sequence of LexA (CTGTN_8_ACAG) (Walker 1984) in the DT40 genome, iChIP-Seq data obtained from Non-KI(B) were used to eliminate genomic regions detected due to such off-target binding (Step 1 in Figure 2). We identified 2,383 peak positions with read numbers more than 2-fold higher in KI(B) than in Non-KI(B), and considered these as potential interacting genomic regions. Because the top 5% peaks (119 peaks) had >7-fold enrichment (Step 1 in Figure 2), we arbitrarily set 7-fold as the threshold for extraction of genomic regions that interact with the *Pax5* promoter region with high frequency. As shown in Step 2 in Figure 2, 105 peaks passed this criterion (>7-fold), from among 2,325 peaks (>2-fold) in the biological replicate. Comparing the 119 (Step 1) and 105 (Step 2) peaks, we identified 34 peaks as reproducibly passing the criterion (Step 3 in Figure 2 and Supplementary Table S1). Among the 34 peaks, 1 peak was the LexA BE-inserted *Pax5* promoter region (Supplementary Table S1) and the other 33 were considered as candidate genomic regions that physically interact with the *Pax5* promoter region.

In this study, we filtered iChIP-Seq data on the basis of the criterion “greater than 7-fold” to extract genomic regions that bind to the *Pax5* promoter region with high frequency. In this regard, more permissive criteria would increase the number of potentially interacting genomic regions. In fact, the criterion “more than 2-fold” extracted 680 common peaks between the 2,383 peaks (Step 1) and 2,325 peaks (Step 2). However, in this case, it might be more difficult to confidently evaluate whether the detected peaks reflect physiological interactions or artificial signals. Therefore, hereafter we focused on the 33 peaks passing the more stringent criterion.

**Figure 2.**
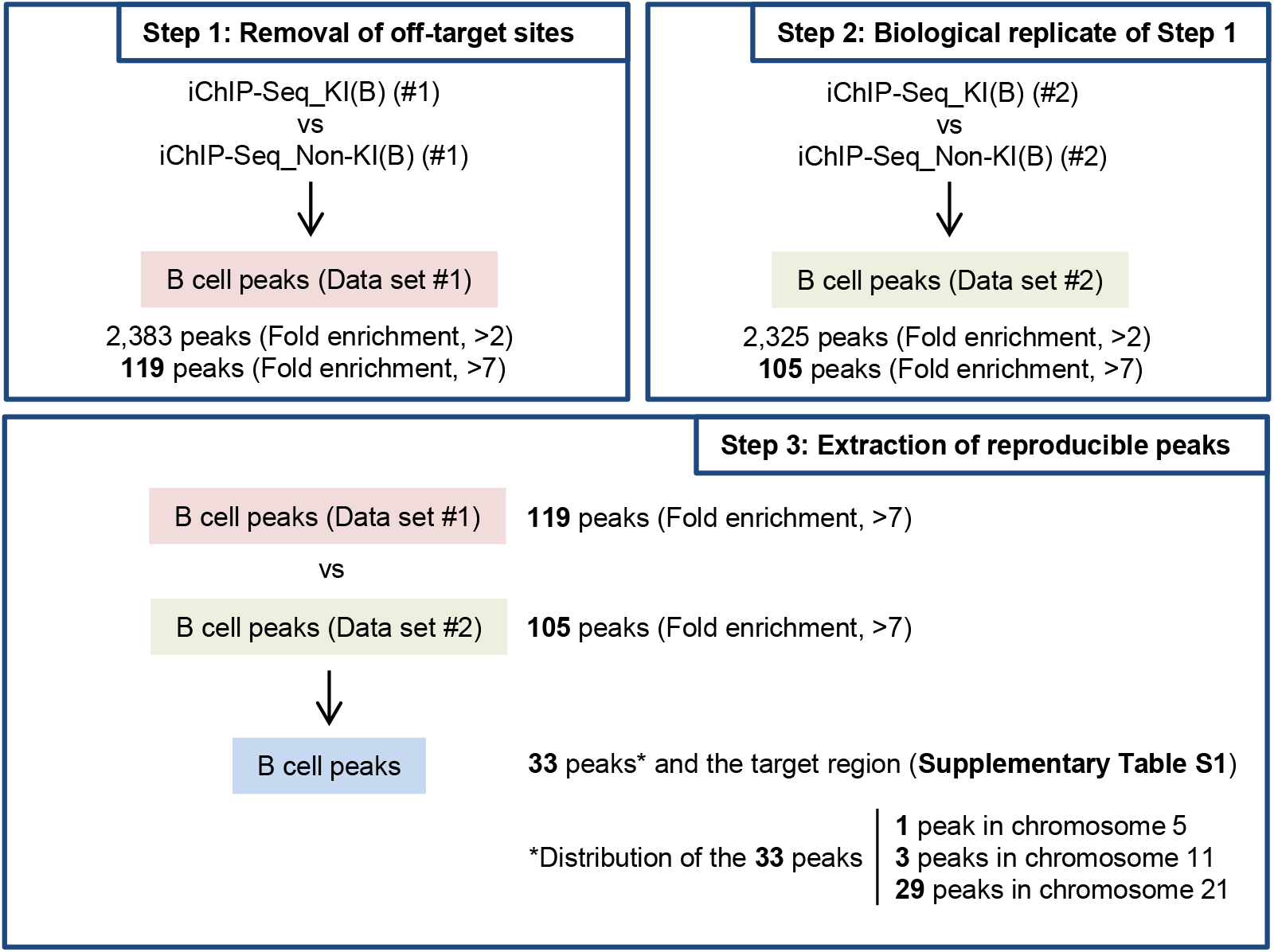
Extraction of genomic regions interacting with the *Pax5* promoter region in DT40. (**Step 1**) Removal of off-target binding sites. iChIP-Seq data were compared between KI(B) and Non-KI(B) (negative control) using Model-based Analysis of ChIP-Seq (MACS) to eliminate off-target binding sites. (**Step 2**) Analysis of another biological replicate of Step 1. (**Step 3**) Identification of genomic regions that interact with the *Pax5* promoter region. The genomic regions commonly detected in Steps 1 and 2 represent candidate genomic regions that physically interact with the *Pax5* promoter region.

### Confirmation of physical chromosomal interactions by *in vitro* enChIP

The chromosomal interactions identified by iChIP-Seq (Figure 2) could include artificial ones caused by insertion of LexA BE. Therefore, it was necessary to confirm the identified chromosomal interactions by another independent method in intact DT40 cells. To this end, we utilized *in vitro* enChIP, an updated form of conventional enChIP (Fujita and Fujii 2014a; Fujita et al. 2016a). In *in vitro* enChIP, recombinant molecules [e.g., recombinant CRISPR ribonucleoproteins (RNPs)] are used for *in vitro* locus-tagging rather than in-cell locus-tagging (Figure 3A). Because intact cells can be utilized in this *in vitro* system, it is unnecessary to consider disruption of physiological chromosomal conformation and potential side-effects caused by in-cell locus-tagging. In *in vitro* enChIP using CRISPR RNPs (Figure 3A), chromosomal conformation in intact DT40 was fixed by formaldehyde crosslinking, and chromatin DNA was fragmented by sonication. The *Pax5* promoter was captured by CRISPR RNPs and isolated from a mixture of the fragmented chromatin by affinity purification. NGS analysis of the isolated material then revealed the genomic regions that physically interact with the *Pax5* promoter.

We designed a guide RNA (Pax5 gRNA) recognizing a 23 bp target site that is 0.1 kb upstream from the TSS of *Pax5* exon 1A (Figure 3B and Supplementary Figure S1); the recognized DNA sequence exists only in the target site, i.e., nowhere else in the chicken genome. We performed *in vitro* enChIP with Pax5 gRNA to specifically isolate the *Pax5* promoter region from intact DT40. Isolation of the *Pax5* promoter region was confirmed by NGS analysis (*in vitro* enChIP-Seq) (Figure 3C) and PCR (Supplementary Figure S1). Next, we examined whether the 33 peaks identified by iChIP-Seq (Step 3 in Figure 2) were also observed by *in vitro* enChIP-Seq (Figure 3D). Based on visual confirmation in Integrative Genomics Viewer (IGV), a high-performance visualization tool, approximately half of the peaks (14 peaks) were also observed by *in vitro* enChIP-Seq in the presence of Pax5 gRNA but not in the absence of gRNA (Table 1 and Supplementary Table S1); representative results are shown in Figure 4 and Supplementary Figures S2 and S3. CRISPR binds to DNA sequences similar to the target sequence, a phenomenon known as off-target binding (Wu et al. 2014; Cencic et al. 2014; O’Geen et al. 2015; Tsai et al. 2015). However, potential off-target binding sites were not found in the 14 identified genomic regions (Supplementary Figure S4). Thus, the genomic regions independently confirmed by *in vitro* enChIP-Seq (Table 1) can be considered as those that physically interact with the *Pax5* promoter region in DT40 cells. These results show that, in addition to iChIP-microarray (McCullagh et al. 2010), iChIP-Seq (and *in vitro* enChIP-Seq) would be a useful tool for non-biased identification of physical chromosomal interactions.

**Figure 3.**
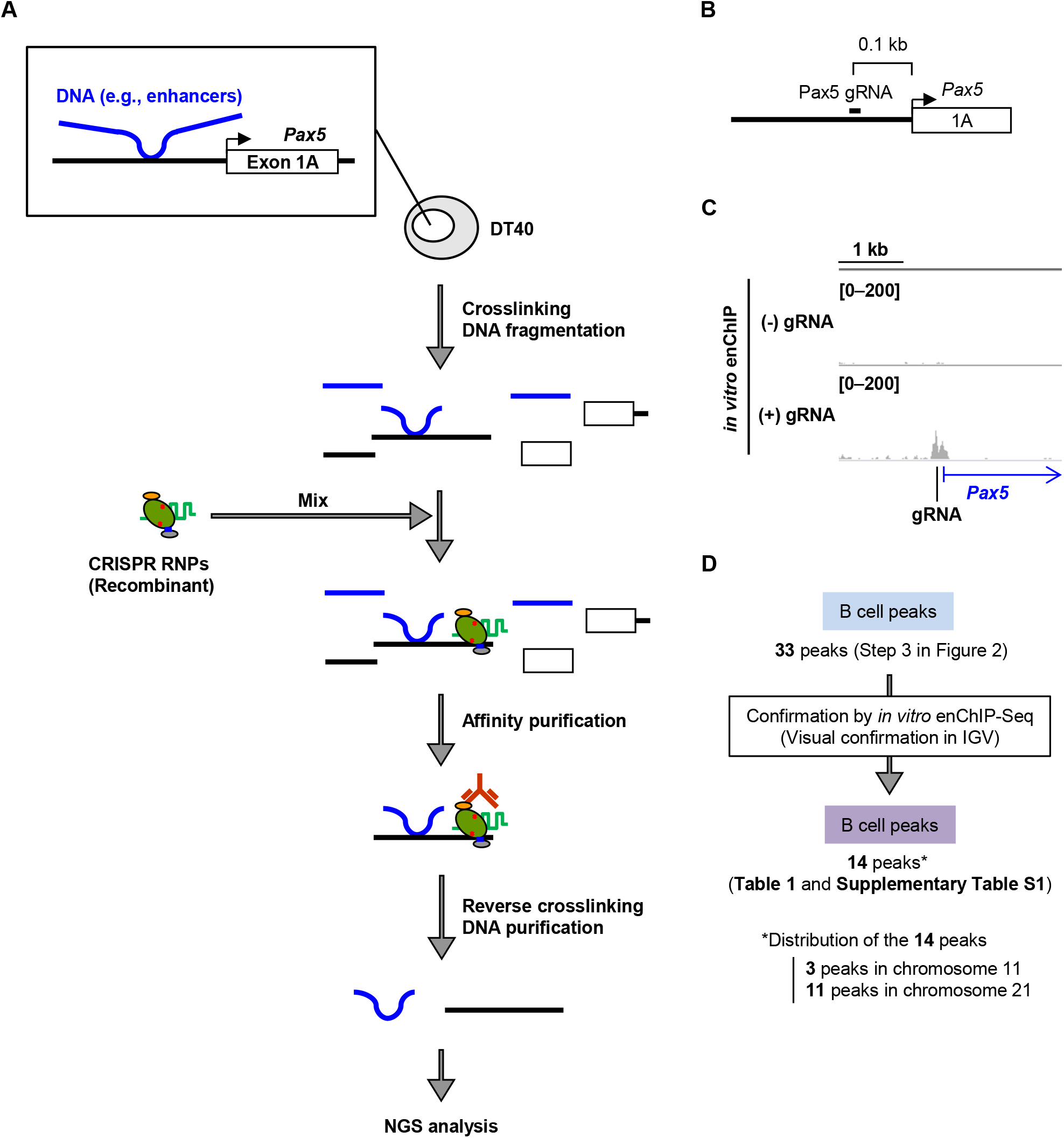
*in vitro* enChIP-Seq for confirmation of the results of iChIP-Seq. (**A**) Intact DT40 cells were crosslinked with formaldehyde, and chromatin DNA was fragmented by sonication. Recombinant CRISPR ribonucleoproteins (RNPs), which consist of 3xFLAG-dCas9-Dock and gRNA targeting the *Pax5* promoter region, were mixed with the fragmented chromatin DNA to capture the target region. After affinity purification with anti-FLAG antibody, reversal of crosslinking, and DNA purification, the DNA was subjected to NGS analysis. (**B**) Target position of *Pax5* gRNA. (**C**) NGS peak images around the *Pax5* promoter region. NGS data from *in vitro* enChIP-Seq were visualized in IGV. The vertical viewing range (*y*-axis shown as Scale) was set at 0–200 based on the magnitude of the noise peaks. (**D**) Confirmation of the results of iChIP-Seq. The peak positions identified by iChIP-Seq were confirmed by *in vitro* enChIP-Seq in IGV.

Interestingly, most of the identified genomic regions were localized in chromosome 21 (Table 1 and Supplementary Table S1). Because those regions were spread equally in chromosome 21 (Supplementary Figure S5), the entire chromosome 21 might be proximal to the *Pax5* gene (or chromosome Z on which the *Pax5* gene is located) in the nucleus of DT40 cells. On the other hand, some peaks identified by iChIP-Seq were not confirmed by *in vitro* enChIP-Seq. In this regard, insertion of LexA BE might partially change the chromosomal conformation around the insertion site and organize artificial chromosomal interactions, which would not be confirmed in intact DT40 by *in vitro* enChIP. Alternatively, *in vitro* enChIP might fail to confirm some *bona fide* chromosomal interactions identified by iChIP-Seq. Because the insertion site of LexA BE is 0.2 kb upstream from the target site of the *Pax5* gRNA (Supplementary Figure S6), iChIP can capture chromosomal interactions organized in the more upstream region of the *Pax5* promoter, whereas *in vitro* enChIP might fail to confirm such chromosomal interactions.

**Figure 4.**
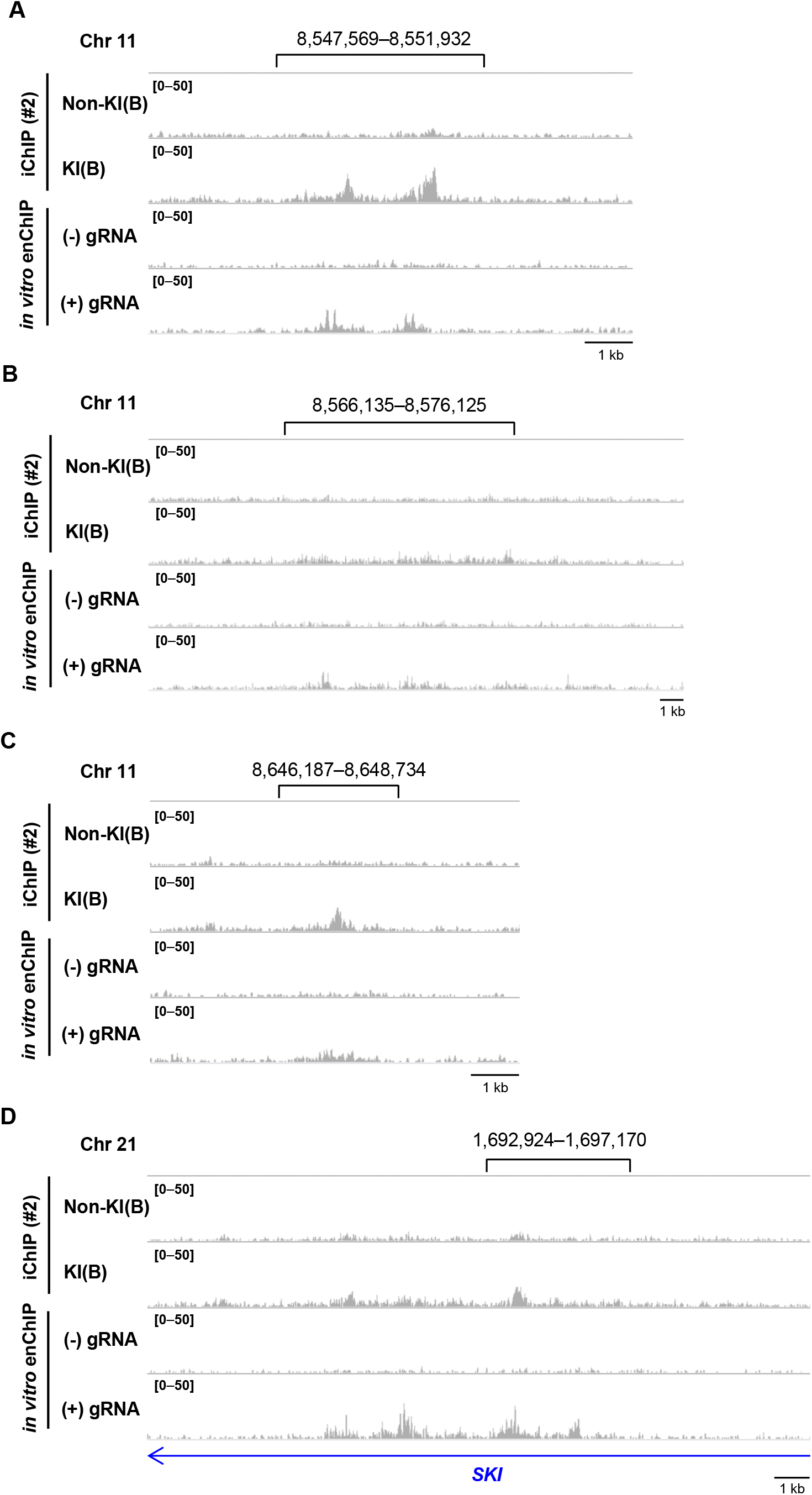
Genomic regions that physically interact with the *Pax5* promoter region. (**A– D**) iChIP-Seq data [#2, Non-KI(B) and KI(B)] and *in vitro* enChIP-Seq data (with or without Pax5 gRNA) were displayed in IGV. Representative regions in chromosome 11 (**A–C**) and chromosome 21 (**D**) are shown. The vertical viewing range (*y*-axis shown as Scale) was set at 1–50 based on the noise peaks. The same loci in the iChIP-Seq data [#1, Non-KI(B) and KI(B)] are shown in Supplementary Figure S2.

**Table 1.**
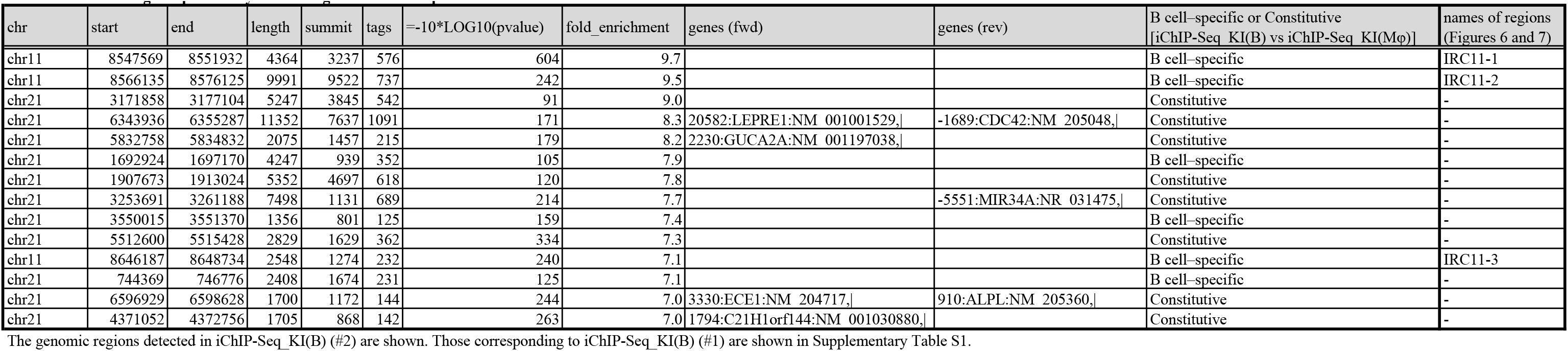
Genomic regions potentially interacting with the *Pax5* promoter in DT40

| chr | start | end | length | summit | tags | =-10*LOG10(pvalue) | fold_enrichment | genes (fwd) | genes (rev) | B cell-specific or Constitutive<br>[iChIP-Seq_KI(B) vs iChIP-Seq_KI(Mφ)] | names of regions<br>(Figures 6 and 7) |
| --- | --- | --- | --- | --- | --- | --- | --- | --- | --- | --- | --- |
| chr11 | 8547569 | 8551932 | 4364 | 3237 | 576 | 604 | 9.7 |  |  | B cell-specific | IRC11-1 |
| chr11 | 8566135 | 8576125 | 9991 | 9522 | 737 | 242 | 9.5 |  |  | B cell-specific | IRC11-2 |
| chr21 | 3171858 | 3177104 | 5247 | 3845 | 542 | 91 | 9.0 |  |  | Constitutive | - |
| chr21 | 6343936 | 6355287 | 11352 | 7637 | 1091 | 171 | 8.3 | 20582:LEPRE1:NM_001001529,\| | -1689:CDC42:NM_205048,\| | Constitutive | - |
| chr21 | 5832758 | 5834832 | 2075 | 1457 | 215 | 179 | 8.2 | 2230:GUCA2A:NM_001197038,\| |  | Constitutive | - |
| chr21 | 1692924 | 1697170 | 4247 | 939 | 352 | 105 | 7.9 |  |  | B cell-specific | - |
| chr21 | 1907673 | 1913024 | 5352 | 4697 | 618 | 120 | 7.8 |  |  | Constitutive | - |
| chr21 | 3253691 | 3261188 | 7498 | 1131 | 689 | 214 | 7.7 |  | -5551:MIR34A:NR_031475,\| | Constitutive | - |
| chr21 | 3550015 | 3551370 | 1356 | 801 | 125 | 159 | 7.4 |  |  | B cell-specific | - |
| chr21 | 5512600 | 5515428 | 2829 | 1629 | 362 | 334 | 7.3 |  |  | Constitutive | - |
| chr11 | 8646187 | 8648734 | 2548 | 1274 | 232 | 240 | 7.1 |  |  | B cell-specific | IRC11-3 |
| chr21 | 744369 | 746776 | 2408 | 1674 | 231 | 125 | 7.1 |  |  | B cell-specific | - |
| chr21 | 6596929 | 6598628 | 1700 | 1172 | 144 | 244 | 7.0 | 3330:ECE1:NM_204717,\| | 910:ALPL:NM_205360,\| | Constitutive | - |
| chr21 | 4371052 | 4372756 | 1705 | 868 | 142 | 263 | 7.0 | 1794:C21H1orf144:NM_001030880,\| |  | Constitutive | - |

The genomic regions detected in iChIP-Seq\_KI(B) (#2) are shown. Those corresponding to iChIP-Seq\_KI(B) (#1) are shown in Supplementary Table S1.

### Identification of genomic regions that physically interact with the *Pax5* promoter region in a B cell–specific manner

The chromosomal interactions identified above might be organized in a B cell–specific manner or occur constitutively in different cell types. Therefore, we next examined whether the 14 interacting genomic regions (Table 1) could be detected in NGS data of iChIP-Seq of KI(MΦ), which are KI(B) trans-differentiated into macrophage-like cells by expression of chicken C/EBPβ (Fujita et al. 2015a). KI(MΦ) expresses M-CSFR, a macrophage marker, but neither Pax5 nor AID, another B-cell marker (Figure 5A) (Fujita et al. 2015a). As shown in Figure 5B, the *Pax5* promoter region was isolated from KI(MΦ) by iChIP. Visual comparison in IGV revealed that three peaks in chromosome 11 and three peaks in chromosome 21 (total six peaks) were observed in a B cell–specific manner, whereas the other eight peaks were constitutively observed both in the B-cell and the macrophage-like cell lines (Table 1); the three B-specific peaks in chromosome 11 and two constitutive peaks in chromosome 21 are shown as representatives in Figure 5D–F and Supplementary Figure S7, respectively. Thus, by comparing iChIP-Seq data, we were able to identify genomic regions that interact with the *Pax5* promoter region in a B cell–specific manner.

We also attempted *in vitro* enChIP-Seq with the macrophage-like cell line DT40(MΦ), which is DT40 trans-differentiated into a macrophage-like cell by ectopic expression of chicken C/EBPβ (Supplementary Figure S8A–C). However, *in vitro* enChIP with Pax5 gRNA failed to isolate the *Pax5* promoter region from DT40(MΦ) (Supplementary Figure S8D), suggesting that the CRISPR RNP was unable to access the gRNA target site. Because *Pax5* transcription is silenced in DT40(MΦ) (Supplementary Figure S8B), the *Pax5* promoter might be heterochromatinized. Alternatively, effector molecules, such as transcriptional repressors, might occupy the gRNA target site, which would block access by the CRISPR RNP.

**Figure 5.**
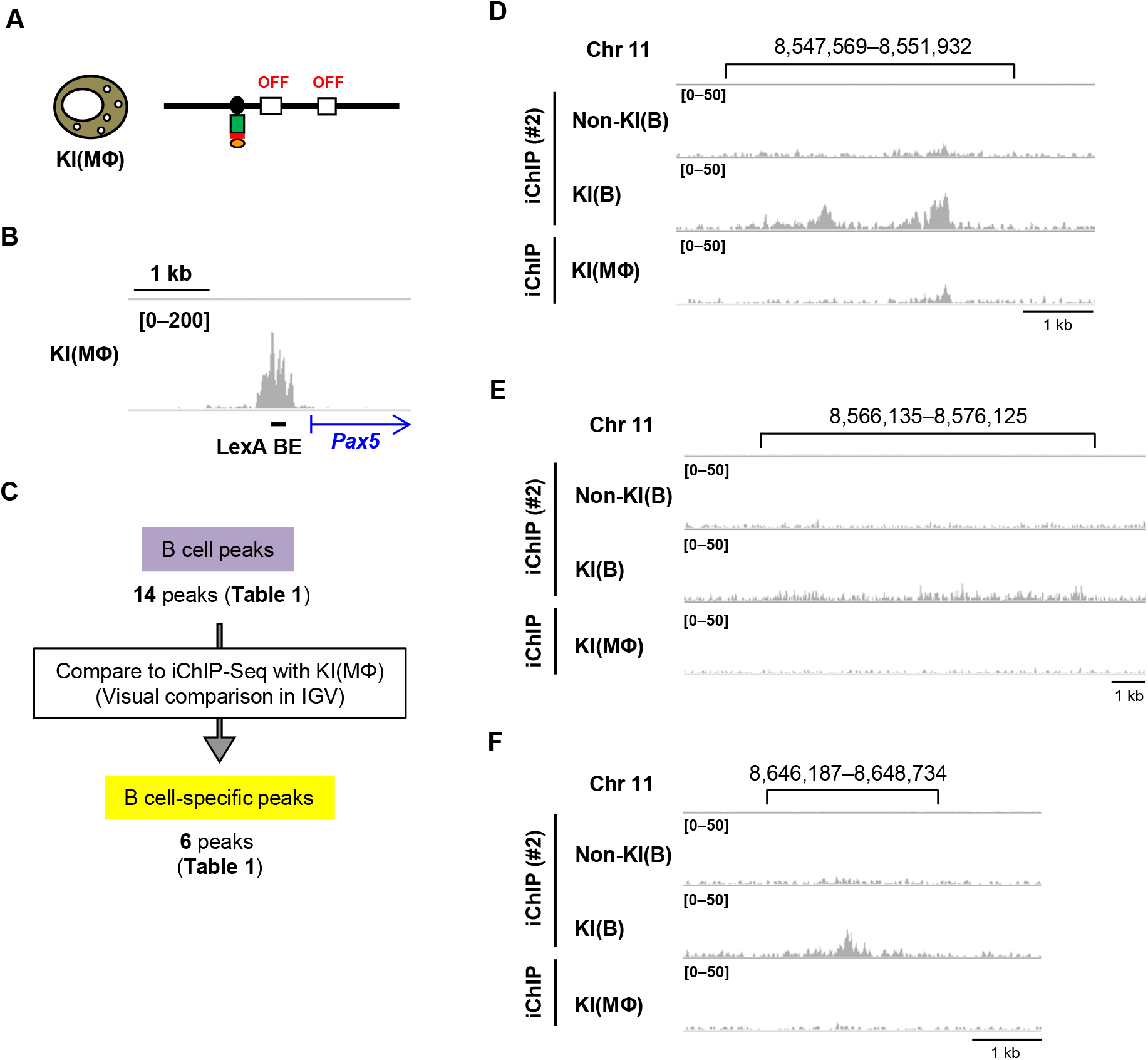
Extraction of genomic regions interacting with the *Pax5* promoter region in a B cell–specific manner. (**A**) In KI(MΦ), which is KI(B) trans-differentiated into a macrophage-like cell, *Pax5* gene is not transcribed. (**B**) Image of NGS peaks around the *Pax5* promoter region. NGS data from iChIP-Seq of KI(MΦ) were visualized in IGV. The vertical viewing range (*y*-axis shown as Scale) was set at 0–200 based on the magnitude of the noise peaks. (**C**) Extraction of genomic regions interacting with the *Pax5* promoter region in a B cell–specific manner. Peak positions identified by iChIP-Seq with KI(B) were compared with the results of iChIP-Seq with KI(MΦ) in IGV. (**D–F**) Representative genomic regions observed in a B cell–specific manner. The images from iChIP-Seq (#2) in Figure 4A–C are also shown here for comparison with those of iChIP-Seq of KI(MΦ).

### Regulation of expression of *Pax5* by a physical interaction between genomic regions

The identified genomic regions (Table 1) might include transcriptional regulatory regions that control *Pax5* transcription through chromosomal interactions. To examine this possibility, we used CRISPR-mediated genome editing to delete genomic regions that B cell–specifically interacted with the *Pax5* promoter region (Jinek et al. 2012; Cong et al. 2013). We chose the three regions in chromosome 11 for locus deletion because they are within 100 kb of each other, and it was therefore feasible to delete all of them at once (Figure 6A), and two of those regions are highly ranked in Table 1. We refer to these three regions (Chr11: 8,547,569– 8,551,932; Chr11: 8,566,135–8,576,125; and Chr11: 8,646,187–8,648,734) as Interacting Region in Chromosome 11 No. 1 (IRC11-1), IRC11-2, and IRC11-3, respectively (Table 1). We constructed plasmids for expression of single guide RNAs (sgRNAs) targeting each end of those genomic regions (Supplementary Figure S9A–C) and co-transfected them with a Cas9 expression plasmid to delete the target genomic regions (Supplementary Figure S9). We were able to delete all three regions (100 kb) in one allele in DT40 (Figure 6B and Supplementary Figure S9D and S10). In the resultant cells (Clone 100k), the transcript levels of the *Pax5* gene were not changed (Figure 6C and D). Next, we deleted each interacting genomic region (IRC11-1, IRC11-2, or IRC11-3) in the other allele in Clone 100k (Figure 6B and Supplementary Figures S9D and S10). Additional deletion of IRC11-2 or IRC11-3 did not have any effects on *Pax5* transcription, whereas deletion of IRC11-1 moderately but significantly decreased transcription of *Pax5* (Figure 6C and D). In DT40, *Pax5* was transcribed comparably from the exons 1A and 1B (Fujita and Fujii 2011b). Deletion of IRC11-1 decreased transcription from exon 1A, but not 1B (Figure 6C and D). To further confirm the physiological importance of IRC11-1 for *Pax5* transcription, we deleted only this region from both alleles in DT40 (Figure 6B and Supplementary Figures S9E and S10). The resultant cells (Clone IRC11-1) also exhibited reduced *Pax5* transcription from the exon 1A (Figure 6C and D). Thus, the decrease in levels of *Pax5* transcription in two independently established cell lines (Clone 100k_IRC11-1 and Clone IRC11-1) suggested that IRC11-1 is involved in transcriptional regulation of the *Pax5* gene, acting as an enhancer via inter-chromosomal interaction.

**Figure 6.**
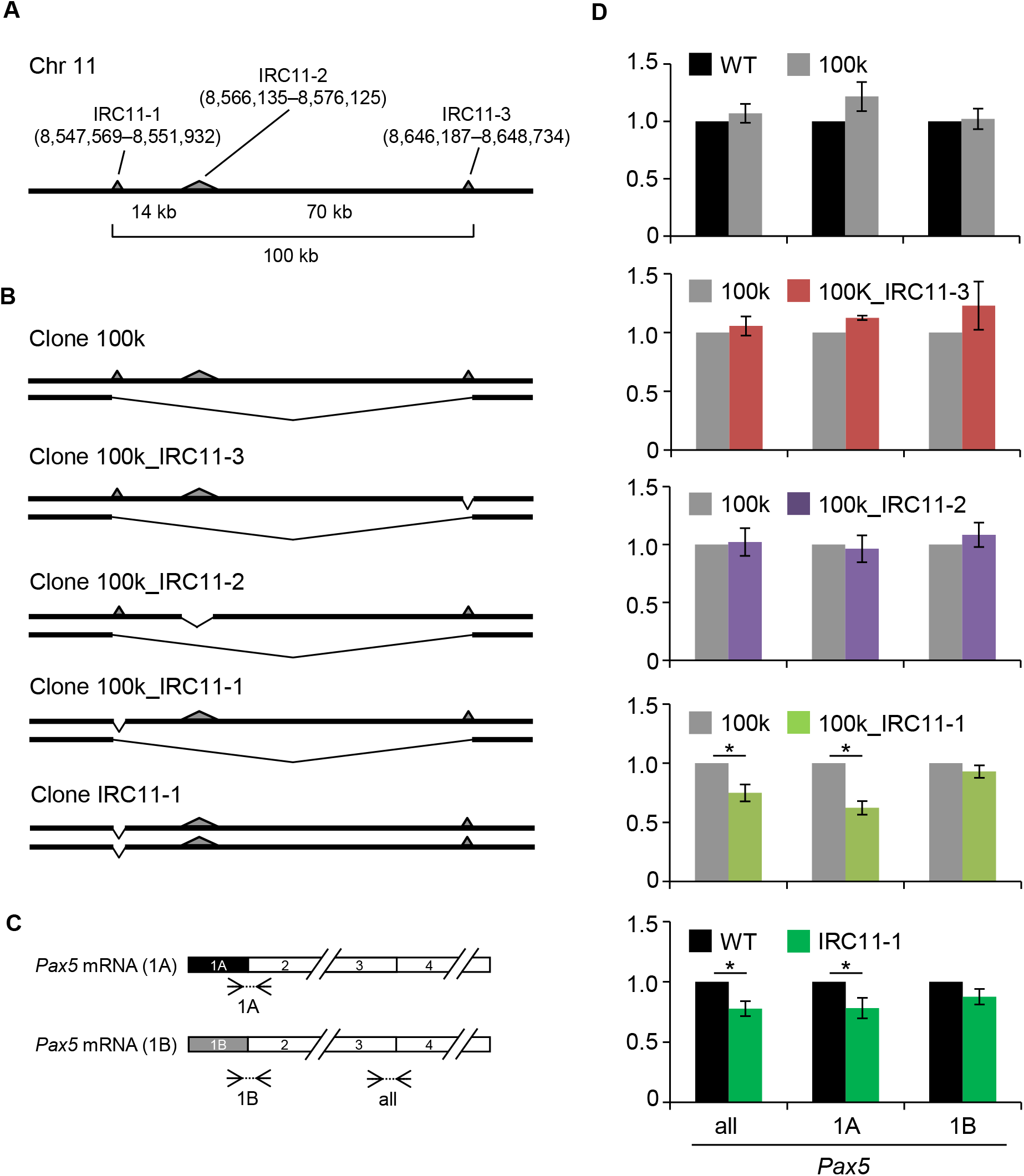
A genomic region interacting with the *Pax5* promoter is involved in transcriptional regulation of the *Pax5* gene. (**A**) Schematic depiction of the loci in chromosome 11 that were identified as interacting with the *Pax5* promoter region in a B cell– specific manner. One allele is shown. The interacting regions are shown as gray triangles. (**B**) CRISPR-mediated knock-out of the interacting regions. (**C**) Primer sets for evaluation of the amounts of *Pax5* mRNA. (**D**) Expression levels of the *Pax5* gene in the knock-out cells. Expression levels of *Pax5* were normalized to those of *GAPDH*, and the mRNA levels in the control cells were defined as 1 [mean ± s.e.m., n = 3 (two upper graphs), n = 6 (others)]. *: t-test p value <0.05. all: mRNA transcribed from both the *Pax5* exons 1A and 1B, 1A: mRNA transcribed from *Pax5* exon 1A, 1B: mRNA transcribed from *Pax5* exon 1B.

Active enhancers are marked by enrichment of histone H3 lysine 4 mono-methylation (H3K4me1) and histone H3 lysine 27 acetylation (H3K27ac) (Heintzman et al. 2007, 2009; Creyghton et al. 2010; Kimura 2013). We therefore investigated whether IRC11-1 is marked by these histone modifications. Because two peaks were observed in IRC11-1 by iChIP-Seq analyses (Figure 4A and Supplementary Figure S2A), we examined these histone modifications at both positions. ChIP assays clearly showed that these histone modifications were enriched at both positions in IRC11-1 but not in an irrelevant genomic region in chromosome 2 (Figure 7), suggesting that IRC11-1 functions as a distal enhancer for *Pax5* transcription. Enrichment of the active enhancer marks was also observed in IRC11-3, whereas only H3K4me1 was enriched in IRC11-2 (Figure 7). Therefore, IRC11-3 might be involved in transcriptional regulation of genes other than *Pax5*.

Deletion of the 100 kb region including IRC11-1 in one allele did not have any effects on *Pax5* transcription (Clone 100k in Figure 6D). Because chromosome Z, which contains *Pax5*, is a single-copy chromosome in DT40 (Sonoda et al. 1998; Chang and Delany 2004), IRC11-1 in each allele may be sufficient for transcription of the single-copy *Pax5* gene. In addition, deletion of IRC11-1 significantly but only partially down-regulated *Pax5* transcription, suggesting that it plays a limited role in *Pax5* transcription. In IRC11-1, two subregions, which are marked by H3K4me1 and H3K27ac, interacted strongly with the *Pax5* promoter (two peak positions in Figure 4A and Supplementary Figure S2A), suggesting that each subregion might work independently or collaboratively to regulate *Pax5* transcription from exon 1A. Future work should seek to elucidate the mechanistic details underlying transcriptional regulation of *Pax5* by IRC11-1.

**Figure 7.**
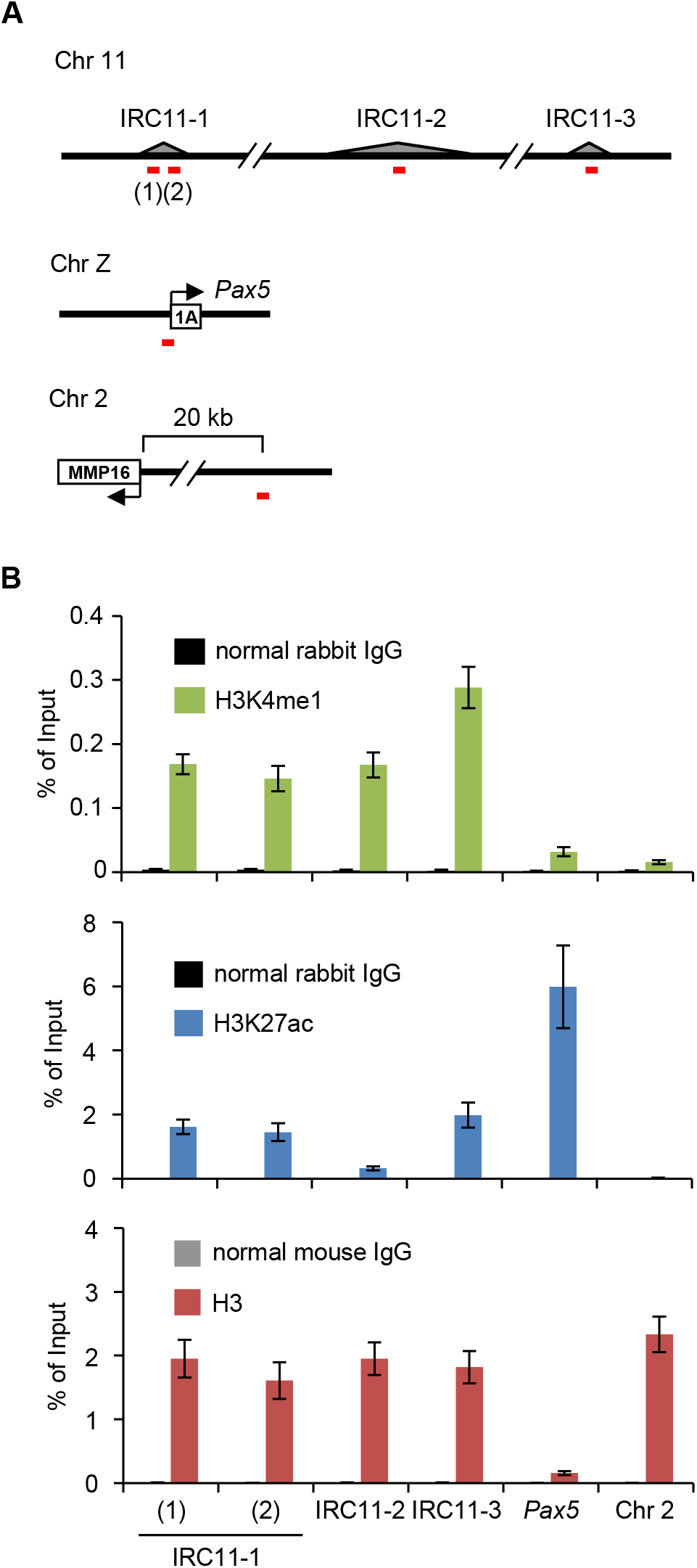
Enrichment of active enhancer marks on the identified genomic regions. (**A**) Positions of primer sets (red lines) used in ChIP assays. (**B**) ChIP assays. DT40 cells were used for ChIP assays with antibodies against H3K4me1 (top), H3K27ac (middle), or H3 (bottom) (means ± s.e.m., n = 3).

### Conclusions and perspectives

In this study, we identified physical chromosomal interactions between the *Pax5* promoter and other genomic regions by locus-specific ChIP in combination with NGS analysis. iChIP-Seq and *in vitro* enChIP-Seq revealed that the *Pax5* promoter binds to multiple genomic regions, in which most regions are localized in chromosome 21 (Figures 1–4). Some of these interactions were organized in a B cell–specific manner (Figure 5). In addition, we showed that deletion of an interacting genomic region in chromosome 11, which is marked by active enhancer histone modifications, decreased transcriptional levels of the *Pax5* gene (Figures 6 and 7), suggesting its physiological involvement in transcriptional regulation of the *Pax5* gene. Our results also indicate that locus-specific ChIP in combination with NGS analysis is a useful tool for performing non-biased searches for physical chromosomal interactions (one-to-many interactions). Thus, this technology could facilitate elucidation of the molecular mechanisms underlying regulation of genome functions, including transcription.

Several methods have been utilized for detection of genome-wide chromosomal interaction. However, observation by only a single method might not accurately reflect physiological chromosomal interactions. In this regard, potential discrepancies have been reported between the results of FISH and those of 3C or its derivatives (Williamson et al. 2014). Therefore, in analysis of chromosomal interactions, it may be necessary to combine several independent methods to eliminate potential contamination of artifactual signals. In this regard, iChIP-Seq and *in vitro* enChIP-Seq could be used as one of several methods. In this study, we used *in vitro* enChIP-Seq to confirm the results of iChIP-Seq. Considering its convenience, *in vitro* enChIP-Seq may be preferable for future identification of chromosomal interactions.

## Materials and Methods

### Plasmids

The Cas9 expression plasmid (Addgene #41815) and chimeric sgRNA expression plasmid (Addgene #41824) were provided by Dr. George Church through Addgene. For construction of the sgRNA expression plasmids, double-stranded DNA (dsDNA) encoding the target sequences were cloned downstream of the U6 promoter in the sgRNA expression plasmid. Alternatively, DNA fragments coding the U6 promoter, target sequence, gRNA scaffold, and termination signal were synthesized and cloned in plasmids by GeneArt gene synthesis services (ThermoFisher Scientific). The target sites of each sgRNA plasmid are shown in Supplementary Figure S9C.

### Cell culture

DT40, Non-KI(B), KI(B), and KI(MΦ) were maintained as described previously (Fujita et al. 2015a).

### iChIP-Seq, *in vitro* enChIP-Seq, and bioinformatics analysis

Non-KI(B), KI(B), and KI(MΦ) (2 × 10^7^ each) were subjected to the iChIP procedure as described previously (Fujita et al. 2015a). DT40 was subjected to the *in vitro* enChIP procedure as described previously (Fujita et al. 2016a). The complex of CRISPR RNA (crRNA) targeting the *Pax5* promoter and trans-activating crRNA (tracrRNA) was used as Pax5 gRNA for *in vitro* enChIP. The gRNA sequences are shown in Supplementary Table S2. Briefly, after fragmentation of chromatin DNA (the average length of fragments was about 2 kbp), the target region was isolated by iChIP or *in vitro* enChIP. After purification of DNA, DNA libraries were prepared using TruSeq ChIP Sample Prep Kit (Illumina); in this preparation step, DNA fragments around 0.4 kbp in length were selectively concentrated. The libraries were subjected to DNA sequencing using the HiSeq platform according to the manufacturer’s protocol. NGS and data analysis were performed at the University of Tokyo as described previously (Yamashita et al. 2011; Seki et al. 2014). Additional information on NGS analysis is provided in Supplementary Table S3. NGS data were mapped onto the reference genome galGal4 using ELAND (Illumina). Narrow peaks of each iChIP-Seq dataset (Steps 1 and 2 in Figure 2) were detected using Model-based Analysis of ChIP-Seq 2 (MACS2, http://liulab.dfci.harvard.edu/MACS/) with default parameters. Images of NGS peaks were generated using IGV (http://software.broadinstitute.org/software/igv/). The accession number of the NGS data is DRA005236.

### Deletion of genomic loci by CRISPR-mediated genome editing

DT40 cells (1 × 10^7^) were transfected with a Cas9 expression plasmid (120 µg), sgRNA expression plasmids (120 µg) targeting each end of a target genomic region, and pEGFP-N3 (0.3 µg, Clontech) by electroporation using a Gene Pulser II (Bio-Rad) at 250 V and 950 µF. One day later, GFP-positive cells were sorted and expanded individually. To confirm targeted locus deletion, genomic DNA was extracted and subjected to genotyping PCR with KOD FX (Toyobo). PCR cycles were as follows: heating at 94°C for 2 min followed by 30 cycles of 98°C for 10 sec, 60°C for 30 sec, and 68°C for 1 min. Primers used for genotyping PCR are shown in Supplementary Table S2.

### RNA extraction and quantitative RT-PCR

Extraction of total RNA and quantitative RT-PCR were performed as described previously (Fujita and Fujii 2012b). Primers used in this experiment are shown in Supplementary Table S2.

### ChIP assays

Antibodies against H3K4me1 (39298, Active Motif), H3K27ac (39134, Active Motif), and histone H3 (MABI0301, Wako) were used. ChIP assays were performed with DT40 cells (2 × 10^6^) and each antibody (3.5 µl for H3K4me1 or 2 µg for the others) as described previously (Fujita and Fujii 2011a). DNA purified using ChIP DNA Clean & Concentrator (Zymo Research) was used as template for real-time PCR with SYBR Select Master Mix (Applied Biosystems) on an Applied Biosystems 7900HT Fast Real-Time PCR System. Primers used in this experiment are shown in Supplementary Table S2.

## Acknowledgements

We thank G.M. Church for providing a plasmid (Addgene plasmid #41815 and #41824) and T. Kikuchi, H. Horiuchi, and M. Tosaka for NGS analysis.

## Data availability

The accession number of the iChIP-Seq and *in vitro* enChIP-Seq data is DRA005236.

